# PK-DB: PharmacoKinetics DataBase for Individualized and Stratified Computational Modeling

**DOI:** 10.1101/760884

**Authors:** Jan Grzegorzewski, Janosch Brandhorst, Dimitra Eleftheriadou, Kathleen Green, Matthias König

## Abstract

A multitude of pharmacokinetics studies have been published. However, due to the lack of an open database, pharmacokinetics data, as well as the corresponding meta-information, have been difficult to access. We present PK-DB (https://pk-db.com), an open database for pharmacokinetics information from clinical trials including pre-clinical research. PK-DB provides curated information on (i) characteristics of studied patient cohorts and subjects (e.g. age, bodyweight, smoking status); (ii) applied interventions (e.g. dosing, substance, route of application); (iii) measured pharmacokinetic time-courses; (iv) pharmacokinetic parameters (e.g. clearance, half-life, area under the curve). Key features are the representation of experimental errors, the normalization of measurement units, annotation of information to biological ontologies, calculation of pharmacokinetic parameters from concentration-time profiles, a workflow for collaborative data curation, strong validation rules on the data, computational access via a REST API as well as human access via a web interface. PK-DB enables meta-analysis based on data from multiple studies and data integration with computational models. A special focus lies on meta-data relevant for individualized and stratified computational modeling with methods like physiologically based pharmacokinetic (PBPK), pharmacokinetic/pharmacodynamic (PK/DB), or population pharmacokinetic (pop PK) modeling.

## INTRODUCTION

The pharmacokinetics (PK) of drugs and medication, i.e., how the body after administration affects substances via absorption, distribution, metabolization, and elimination, are of great interest for medical research and drug development. The main measures in the field are concentration-time profiles and derived PK parameters from these timecourses like half-lifes or clearance rates. These measures strongly depend on the dosage and individual characteristics of the subject or group under investigation. Factors like age, weight, sex, smoking behavior, or disease drive the large inter-individual variability in PK (17) making such meta-data indispensable for research in pharmacokinetics. The study of variability in drug exposure due to these covariates is an important field of research with a long history, generally referred to as population pharmacokinetics (1). Modern approaches go beyond classical population information by accounting for additional factors, for example, for genetic variants (12). This meta-information on subjects in combination with the main measures are the basis for individualized and stratified approaches in drug treatment which will potentially pave the road towards both precision dosing and precision medicine.

A multitude of PK studies have been published but despite the wealth of literature almost none of the data is accessible in a machine-readable format and certainly not with FAIR (findable, accessible, interoperable and reproducible) principles (26) in mind. The lack of transparency and reproducibility (11) in the field is ubiquitous. Currently the only way to retrieve this treasure is by digitizing and curating the pharmacokinetics information from publications. Despite the central role of PK in the medical and pharma field, or perhaps exactly because of that, no open freely accessible database of pharmacokinetics information exists so far. In addition, heterogeneity in the reporting of clinical study designs, pharmacokinetic measures, individual, and population-related meta-information further complicates data reuse and integration. Many studies only report a small fraction of the underlying data, e.g., individual data or prominent PK parameters are missing in most studies and even averaged time-courses are only present in a subset of data.

For computational modeling, meta-analysis, and most methods in machine learning a standardized and machine-readable representation of data is of major importance. PK data could be utilized in many different ways (18, 19, 24) if such a representation and corresponding database would exist. One such application is physiologically based pharmacokinetic modeling (PBPK) which provides a unique opportunity to integrate PK data and parameters from multiple clinical trials into a single model. These models can account for the differences in the study protocol, the dosing, as well as individual, group and population characteristics.

## DESCRIPTION & RESULTS

PK-DB (https://pk-db.com) is an open-source web-accessible database storing comprehensive information on pharmacokinetics studies consisting of PK data, PK parameters, and associated meta-information (see Figure 1 for a general overview).

**Figure 1.**
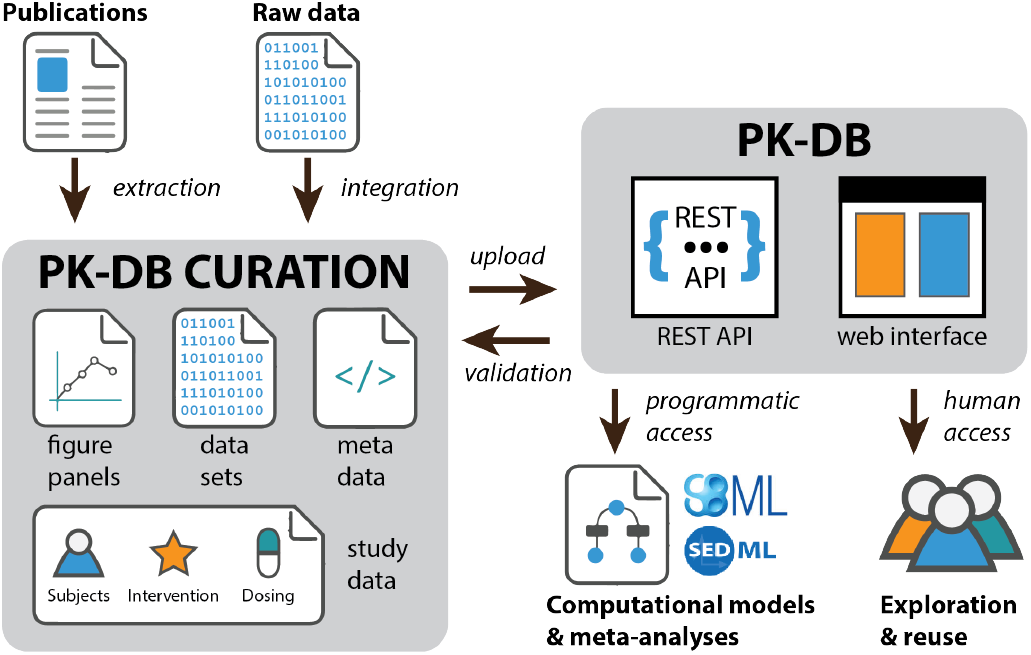
PK-DB overview. Schematic overview of the curation process and interaction with the PK-DB database. Data is either extracted from literature (digitization of figures and tables) or data sets are directly imported (from collaboration partners). Figure panels, data sets, meta-data and study information on subjects, interventions and dosing is curated. All data files and the study information are uploaded via REST endpoints. The curated data is checked against validation rules, data is normalized (e.g., units), and pharmacokinetic parameters are calculated. The uploaded study information can either be programmatically accessed via the REST API or via the web frontend.

### Database statistics

PK-DB provides curated information on (i) characteristics of studied patient cohorts and subjects (e.g. age, bodyweight, smoking status); (ii) applied interventions (e.g. dosing, substance, route of application); (iii) concentration-time curves; and (iv) parameters measured in PK studies (e.g. clearance, half-life, and area under the curve). The focus so far of data curation has been on substances applied in dynamical liver function tests and studies of glucose metabolism.

PK-DB-v0.6.5 (15) consists of 183 studies containing 473 groups, 1808 individuals, 510 interventions, 15790 outputs and 1260 time-courses related to caffeine, glucose, codeine, or paracetamol (see Figure 2 and Supplementary Material 1, Supplementary Material 2, and Supplementary Material 3).

**Figure 2.**
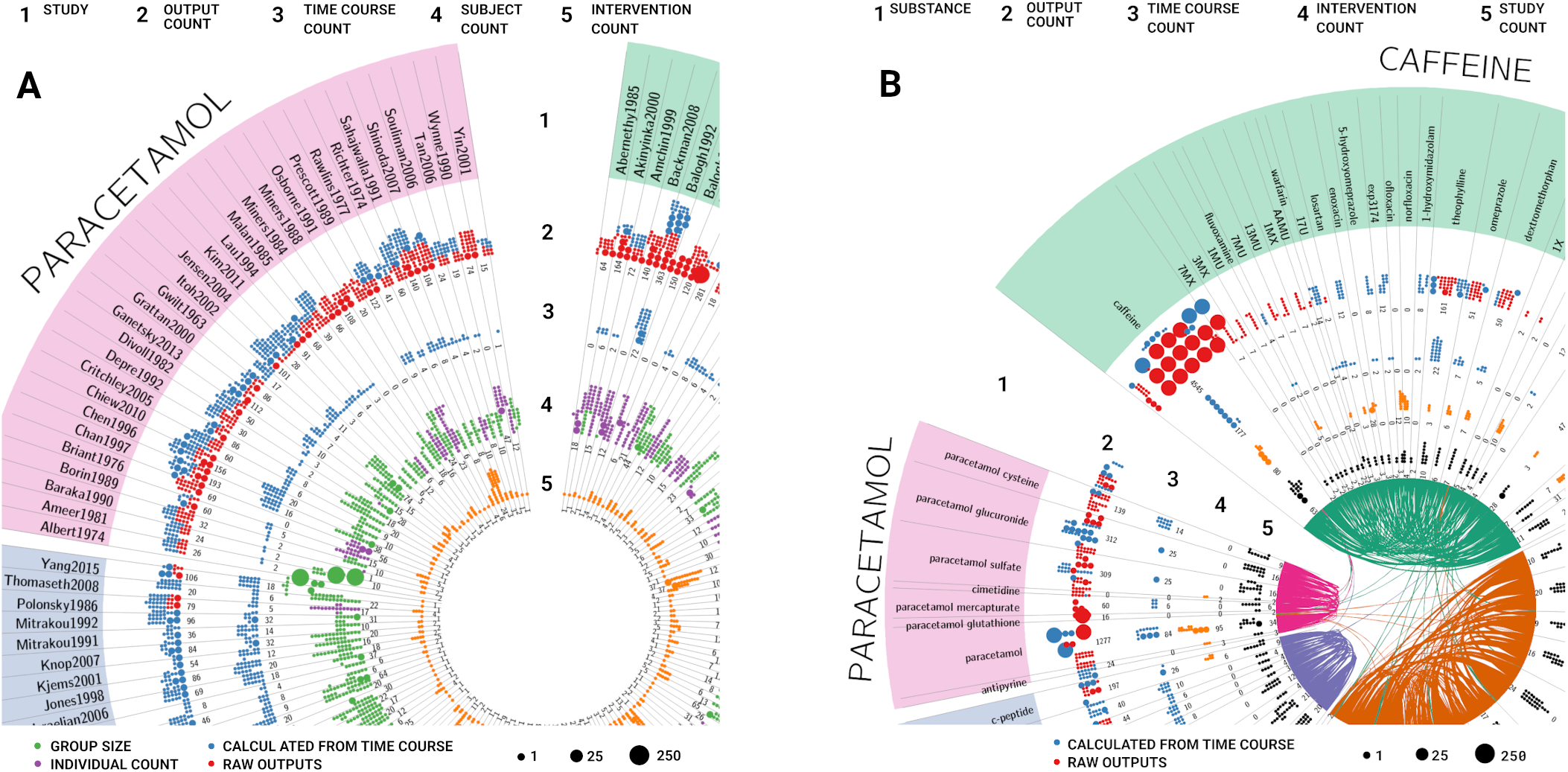
PK-DB content. **(A) Studies.** Overview of the study content in PK-DB. The complete data is available in Supplementary Material 1 and Supplementary Material 3. PK-DB-v0.6.5 (15) consists of 183 studies containing 473 groups, 1808 individuals, 510 interventions, 15790 outputs and 1260 time-courses related to caffeine, glucose, codeine, or paracetamol. The circular plot is structured in stripes and rings, with each stripe representing a single study. In each ring, the counts of different data types are depicted. Dot size corresponds to the number of entries. The rings give an overview of the following information (1) name of the study; (2) number of outputs (PK parameters and other measurements). Red dots represent reported data, blue dots data calculated from time-courses; (3) number of time-courses; (4) number of participants. Purple dots represent participants with individual data, green dots represent participants which are reported as a group; (5) the number of interventions applied to the participants in the study. **(B) Substances** Overview of the substance content in PK-DB. The complete data is available in Supplementary Material 2 and Supplementary Material 3. Substances with very few entries (*<*2 studies) are excluded from the plot. The circular plot is structured in stripes and rings, with each stripe representing a different substance. Substances were clustered in five substance classes (caffeine, glucose, codeine, and paracetamol) by agglomerative clustering of the pair co-occurrence of substances within studies. Classes are labeled according to the most frequent substance within the class. Each co-occurrence of two substances is visualized by a connecting ribbon between the substances in the center. The rings describe the following information for the respective substance (1) name of the substance; (2) number of outputs (PK parameters and other measurements). Red dots represent reported data and blue dots represent data calculated from reported concentration-time profiles. (3) the number of time-courses; (4) number of applied interventions; (5) number of studies in which the substance occurred.

### Design principles

Important features of PK-DB are the representation of experimental errors, the normalization of measurement units, annotation of information to biological ontologies, calculation of pharmacokinetic parameters from concentration-time profiles, a workflow for collaborative data curation, strong validation rules on the data, computational access via a REST API as well as human access via a web interface. Key principles in the design of PK-DB were:

#### Accessibility of data for computational modeling and data science

All data is available via REST endpoints allowing for simple integration of PK-DB data into existing workflows, e.g., for the building of computational models. The major advantage of a REST API as a central access point to the database is that it can be accessed from various clients independent of the programming language. In the following, we present various use cases to demonstrate the usefulness of this approach, e.g., creating an overview of the database content using R and circos (Figure 2), and meta-analyses of multiple studies using Python (Figure 4). The use of PK-DB data is facilitated by annotation of biological and medical concepts to respective ontologies. This enables the integration with additional data sets or computational models based on the semantic information, e.g., substances are annotated to ChEBI (10), and diseases to ncit, hp, doid, and mondo (14, 16, 20, 22). A special focus lies on meta-data for individualized and stratified computational modeling with methods like physiologically based pharmacokinetic (PBPK), pharmacokinetic/pharmacodynamic (PK/DB), or population pharmacokinetic (pop PK) modeling.

#### Extensibility and generalizability

The PK-DB data model is not limited to a specific problem domain but allows simple extensions to other fields and experimental data sets, within the overall area of pharmacokinetics. Examples are extensible types for the group or individual characteristics currently represented in the database. Additional types can easily be added to cover the important information for a given problem domain.

#### Unit and data normalization

A key challenge in using data for computational modeling and data science are non-standardized units coming from different data sets. It requires time-consuming retrieval of this information from the literature and error-prone conversion of units and corresponding data. PK-DB provides a solution to this issue. During upload the data is harmonized, e.g., data is converted between molar units and gram, using thereby the molecular mass of the respective substances based on its ChEBI information (10). In addition, for all information stored the allowed units are defined (actual units must be convertible to these units).

#### Representation of time-course data

The main measures in pharmacokinetics studies are concentration-time curves of the administered substance and its metabolites after biotransformations. These time-courses are crucial for kinetic modeling, e.g., using physiologically based pharmacokinetic (PBPK) or pharmacokinetics/pharmacodynamics (PK/PD) models. Consequently, a central focus was on storing and analyzing such data efficiently.

#### Calculation of pharmacokinetic parameters

PK-DB calculates important secondary PK parameters such as half-life, clearance or volume of distribution from the time-concentration profiles during data upload based on non-compartmental methods (6). Parameters are calculated based on linear regression of the logarithmic concentration values in the exponential decay phase (see example in Figure 3 and Table 1). Non-compartmental methods were chosen for comparison of calculated values with reported PK parameters in the literature, mainly calculated based on non-compartmental methods.

**Table 1.**
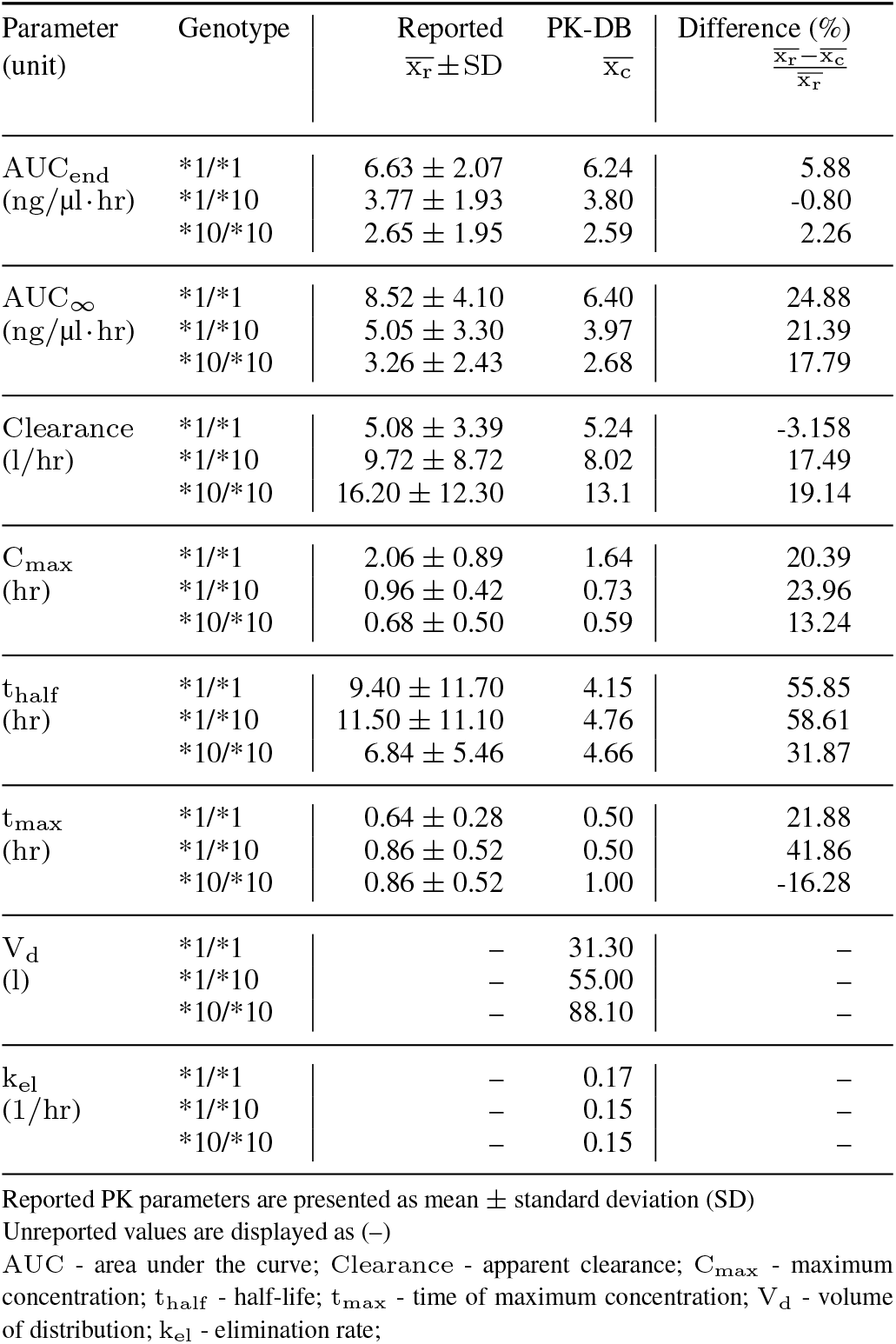
Calculation of pharmacokinetic parameters. Comparison of PK parameters reported in a representative study of codeine (27) with PK parameters calculated from mean concentration-time profiles (see Figure 3). Only data for the groups, no individual data was reported in the study.

**Figure 3.**
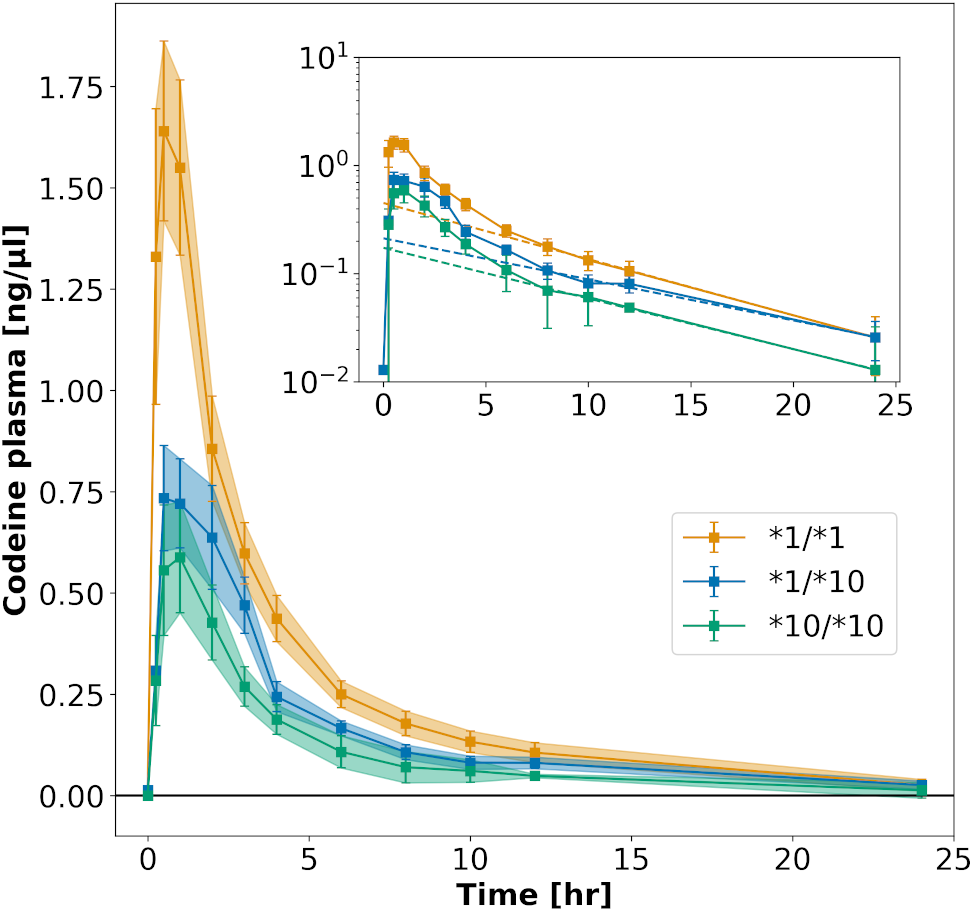
Calculation of pharmacokinetics data from time-courses. Concentration-time courses have been digitized from figures in the literature. This example shows codeine plasma time-courses after the application of codeine to three subgroups with different genotypes (27). Note that concentration as well as time units are automatically normalized. PK parameters are calculated from reported concentration-time profiles using non-compartmental methods, e.g., the apparent clearance of codeine, the half-life (t_half_) of codeine or the volume of distribution (V_d_). The exponential decay is used for the fitting of PK parameters (see inlet). Calculated and reported PK parameters for this example are listed in Table 1. Due to the unavailability of individual participant data in most pharmacokinetics studies, parameters have to be determined on the mean time-concentration curves (averaged over subjects in a given group).

**Figure 4.**
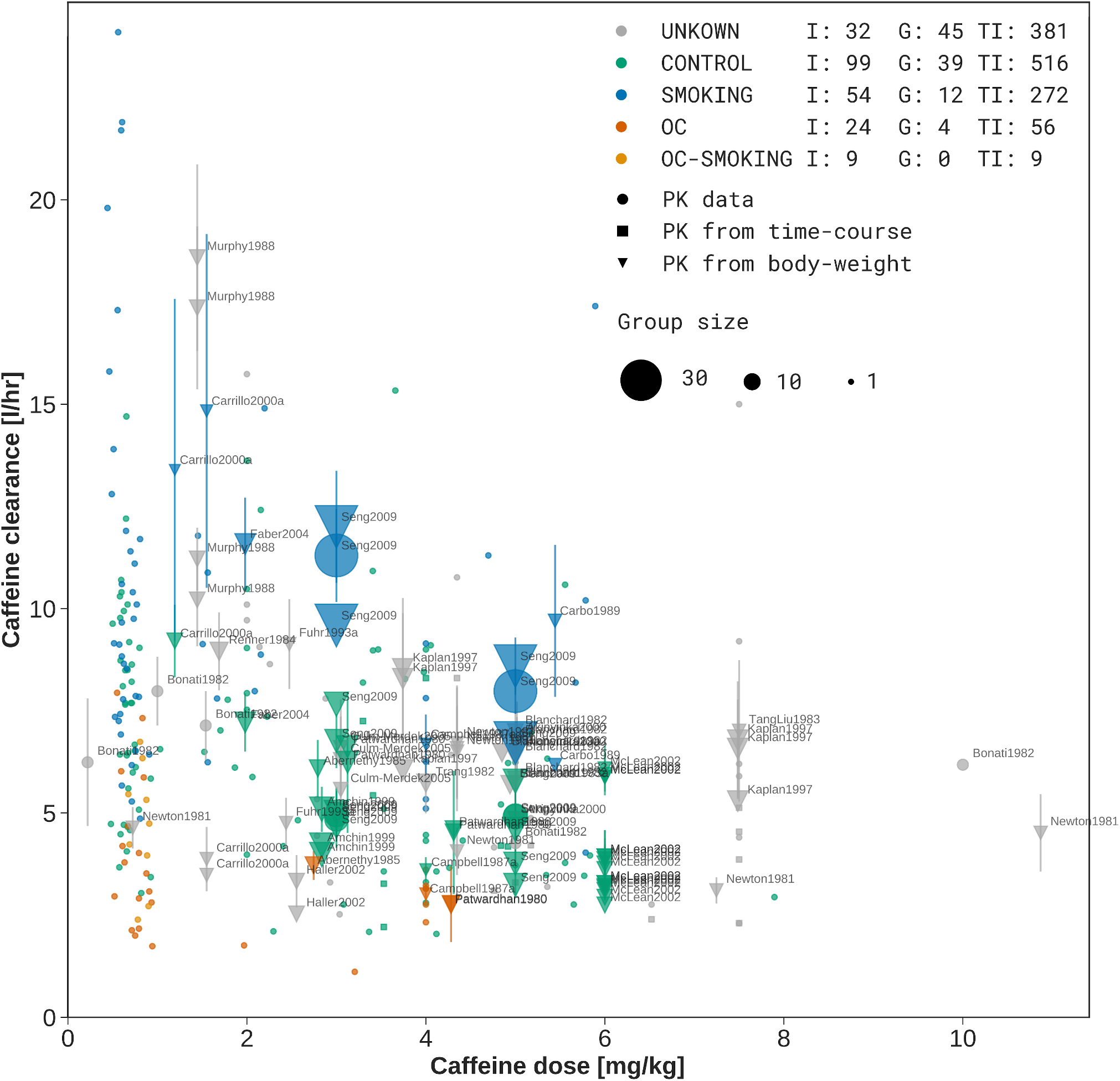
Meta-analysis of caffeine clearance depending on caffeine dose. Caffeine clearance is stratified based on reported smoking and oral contraceptive (OC) use. UNKNOWN (grey) data corresponds to unreported smoking and OC, CONTROL (green) are non-smokers not taking OC, SMOKING (blue) are smokers not taking OC, OC (dark orange) are non-smokers taking OC, and OC-SMOKING (light orange) are smokers taking oral contraceptives. For the stratification groups the number of individuals (I), number of groups (G) and number of total participants (TP) is provided in the legend. Individual and group data is depicted, with group size encoded as dot size. Data points from groups are labeled by the study identifier. Reported PK parameters are depicted as circles, PK parameters calculated from concentration-time profiles as squares, and PK parameters inferred from PK data and reported bodyweights of the participants as triangles (to convert to dose per bodyweight). Typically, dosing is reported in mass units and clearance in a volume per time. Sometimes both values are reported in bodyweight units. Here, all available data is harmonized. Suspicious data from four studies (2, 8, 9, 23), very likely from a single clinical trial, was excluded.

#### Data quality

Strong validation (e.g. of categoricals), minimum relevant information, instance cross-referencing and correct unit-dimensions ensure high quality of the curated data. Non-obvious curation mistakes (or respective errors in the reporting of the data) can be addressed by outlier identification in subsequent meta-analyses.

#### Access rights

PK-DB allows to keep studies privately or only share with certain collaborators. This allows sharing the study during the curation process only with trusted people with a simple option to make the study public. Some information is only accessible by a limited group of users due to copyright issues, e.g., for manually curated studies from the literature the underlying publication can only be made accessible if it is Open Access. A subset of studies is currently private because the underlying raw data from the clinical trial has not been published yet.

### Technology

The PK-DB backend is implemented in Python using the Django framework with Postgres as the underlying database system. For fast, full-text search most data is indexed with Elasticsearch. The provided REST API uses the Django-rest-framework with endpoints accessible from https://pk-db.com/api/. The web frontend (https://pk-db.com) is implemented in JavaScript based on the Vue.js framework interacting with the backend via the REST API. The complete PK-DB stack is distributed as docker-containers. PK-DB is licensed under GNU Lesser General Public License version 3 (LGPL-3.0) with source code available from https://github.com/matthiaskoenig/pkdb.

### Curation workflow

PK-DB provides a collaborative curation interface which simplifies the upload and update of curated study information. A central component is to track all files and curation changes via Git version control. On changes, the files can automatically be uploaded and validated against a development server which provides direct feedback on missing information or curation errors (e.g., units for bodyweights must be convertible to [kg]). A multitude of constraints have been defined as validation rules on the uploaded data instead of having the data model layer too restrictive. These validation rules are constantly updated based on curator feedback. Allowed choices in the data model are based on an internal ontology, which allows to update encodable information without the need to update the database backend.

The typical workflow for extracting data from the literature is depicted in Figure 1. At the beginning of the curation process, a body of literature is selected based on literature research for a given problem domain. Subsequently, the relevant (meta-)information is manually extracted from the literature and encoded in a standardized JSON format. Extracted data like concentration-time courses or PK parameters are stored as tabular data in spreadsheets. After finishing the initial curation process, a second curator is checking the data.

Curation is an iterative process involving multiple curators over time. Tracking changes to the curated data is therefore crucial. Instead of implementing such history and change tracking on database level with substantial overhead, we utilize the full set of Git features out of the box to track changes to our files. All curators work hereby on a shared Git repository. Private data can be tracked in separate private Git repositories.

### Calculation of pharmacokinetic parameters

An important part of PK-DB is the automatic calculation of PK parameters from the reported concentration-time curves based on non-compartmental methods (6). Figure 3 and Table 1 illustrate the automatic calculation of PK parameters from concentration-time profiles for an example study. The authors were hereby interested in the influence of specific genetic alleles on the pharmacokinetics of codeine (27). In the study, information was limited to the averaged measures with variation (standard error within group), but no individual subject data was reported.

Calculated parameters are the area under the curve (AUC_end_), the area under the curve extrapolated to infinity (AUC_∞_), the concentration maximum (C_max_), the time at concentration maximum (t_max_), the half-life (t_half_), the elimination rate (k_el_), the clearance (Clearance) and the volume of distribution (V_d_). The calculated values are in good agreement with the reported values (all lie within the reported standard deviations).

Mathematically correct, first the PK parameters should be calculated for each subject individually and subsequently be averaged. Unfortunately, this is not possible if only averaged data is reported. Consequently, as approximation PK parameters are calculated on the averaged time-courses. Due to the often very large interindividual differences in pharmacokinetics the calculated values on average data can be notable different between reported and calculated parameters (Table 1, e.g., t_half_). Even more fundamentally, the description of the data as averages with variations has inherent problems by assuming homogeneity of the data which often is not the case (7). Consequently, we strongly encourage the publication of individual subject data in PK studies.

A further limitation of PK studies is that often only a subset of pharmacokinetics information is reported. In the example displayed in Table 1 (27), the volume of distribution (V_d_) and the elimination rate (k_el_) are not reported, but can be calculated.

### Meta-analysis of caffeine

PK-DB allowed us for the first time to undertake an extensive and systematic analysis of the effect of lifestyle factors like smoking and oral contraceptive use on the clearance of caffeine combining data from multiple studies. For this use case we integrated data from 44 studies, based on programmatic interaction with PK-DB via the REST API. By curating information about the respective patient characteristics (lifestyle factors), the actual interventions performed in the studies (dosing and route), and important information like the errors on the reported data we could gain a unique view on the strong and consistent effect of smoking and oral contraceptive use on the clearance of caffeine. The large variability between studies and individuals could be markedly reduced by accounting for lifestyle information.

Importantly, the meta-analysis allowed us to directly improve the curation status of many studies by easily detecting visible outliers in the data which could in most cases directly be backtracked to curation errors or incorrectly reported data (e.g., incorrect units) which were subsequently corrected in the database.

A positive aspect is that most of the reported studies are consistent. For instance with caffeine, most of the data was in line with each other with a single exception being Stille et al. (23). Here a systematic bias in the data could be observed probably due to an analytic problem. Interestingly, the same data set was published multiple times, overall in four publications all showing the same bias (2, 8, 9, 23).

### Data quality and validation

The integration of data from multiple studies and subsequent meta-analyses is a valuable procedure to identify curation errors which cannot be caught by validation rules alone. The combination of both, the validation rules and the meta-analyses helped to identify errors also in the reporting. In the following, we will give some examples of suspicious reported data detected by meta-analysis: Wang et al. (25) reported incorrect units; Seng et al. (21) calculated volumes per bodyweight incorrectly; In the publication of Carbo et al. (4) participant number 4 has a suspiciously high half-life and participant number 3 a suspiciously high clearance rate. It is unclear if this is a reporting error; In the publication of Beach et al. (3) 9 smokers and 2 non-smokers have suspiciously very high clearance rates, again unclear if this is a reporting error. In the publication of Wu et al. (27) the concentration-time profiles and concentration maxima were reported with incorrect units.

Data validation and data integration via PK-DB allowed us to identify and correct these issues.

## CONCLUSION & DISCUSSION

PK-DB is the first open database for pharmacokinetics data and corresponding meta-information. We provide an important resource which allows storing pharmacokinetics information in a FAIR (findable, accessible, interoperable and reproducible) manner (26). We demonstrate the value of PK-DB via a stratified meta-analysis of pharmacokinetics studies for caffeine curated from literature which allows us to integrate and harmonize pharmacokinetics information from a wide range of studies and sources.

By performing the curation for commonly applied drugs (codeine and paracetamol), for a substance used in liver function tests (caffeine), as well as for glucose we could demonstrate the applicability of PK-DB to a wide range of substances and gain insights into how well data is reported in the various fields.

The reporting of data in the field of pharmacokinetics is very poor despite the main point of the publications being the reporting of the data. Without guidelines on minimal information for studies, it is very difficult to compare studies or integrate data from different sources. Incomplete and poor reporting of data in the field of pharmacokinetics has also been reported by others (5, 13). As our analysis shows, even basic information, crucial for the interpretation and analyses of PK studies, are not reported in many publications. It is impossible to integrate and reuse such data. For instance, in the case of codeine, often not even the given dose can be retrieved from the publication because it is not clearly reported which substance was administered (codeine-sulfate, codeine-phosphate or codeine). Other examples are unreported bodyweights, so that conversions to doses per bodyweight are not possible.

Based on our work we have a set of important suggestions when publishing clinical studies in the field of pharmacokinetics: (i) Publish the actual data in a machine-readable format (e.g., a data table in the supplement); (ii) Publish the actual concentration-time curves, not only derived parameters; (iii) Provide data for individual subjects which is much more informative and allows to calculate all data for individuals and for groups; (iv) Provide minimum information on (individual) patient characteristics which includes basic anthropometric information like age, bodyweight, sex, height, and the subset of important lifestyle factors known to alter pharmacokinetics (e.g. co-medication, oral contraceptive use, smoking status, alcohol consumption or for instance for CYP1A2 substrates like caffeine: methylxanthine consumption/abstinence); (v) Clearly state the study protocol: Which substance was given in which dose, in which route (oral, intravenous), and in what form (tablet, capsule, solution), the more specific the information the better.

We envision that PK-DB will encourage better reporting of pharmacokinetics studies by providing means for data representation and integration and will improve reusability of pharmacokinetics information by providing PK data in a central database, and will facilitate data integration between studies and with computational models.

## Contributions

JG and MK conceived the study, drafted the manuscript, wrote the software, and performed all analyses. JG, MK, DE and KG curated studies for PK-DB. All authors read, corrected and approved the manuscript.

## SUPPLEMENTARY MATERIAL

**Supplementary Material 1** PK-DB study overview

**Supplementary Material 2** PK-DB substance overview

**Supplementary Material 3** PK-DB data overview

