## Supplementary figures and images for "PK-DB: PharmacoKinetics DataBase for Individualized and Stratified Computational Modeling"

### Supplementary Material 1

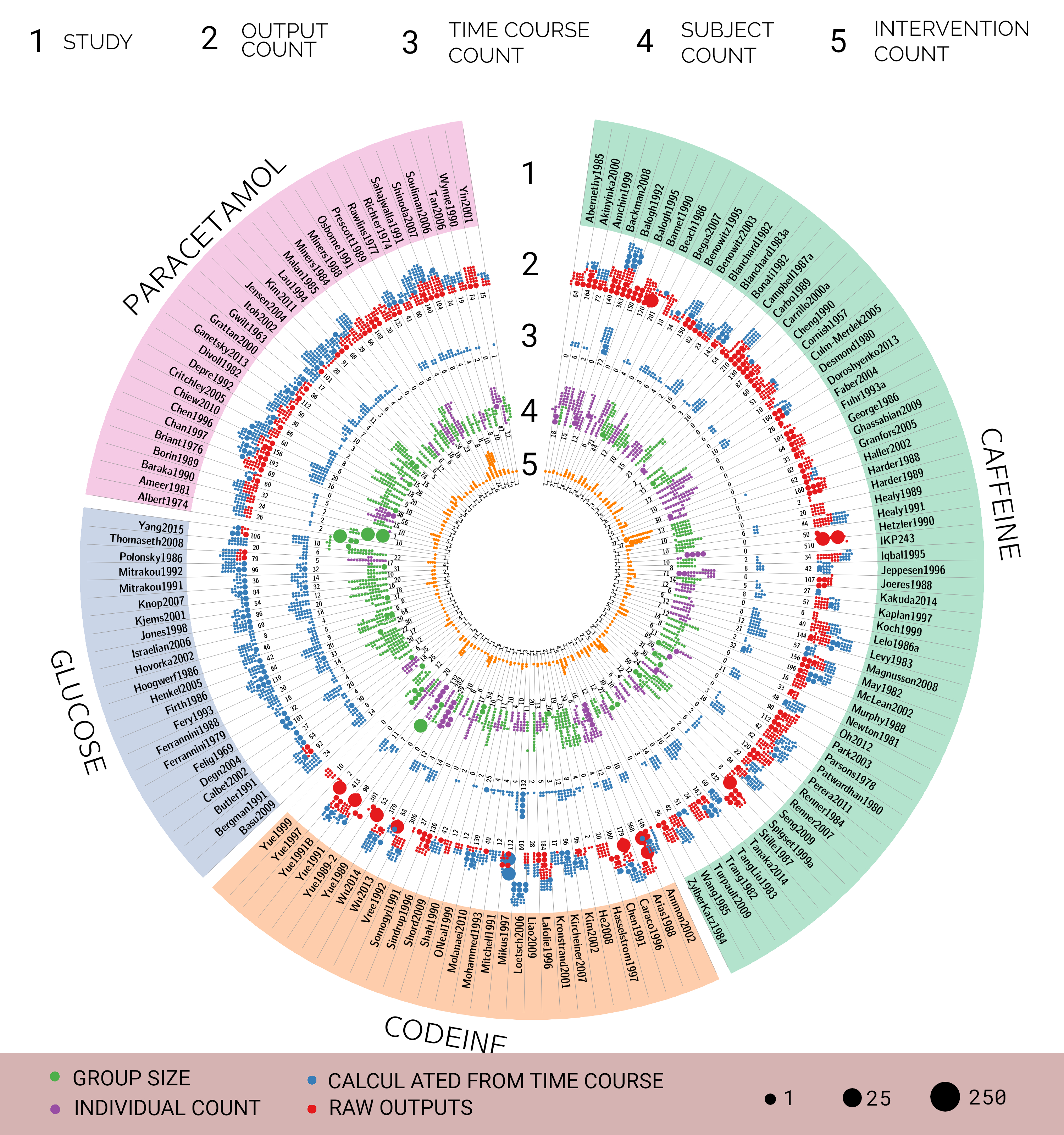

### Supplementary Material 2

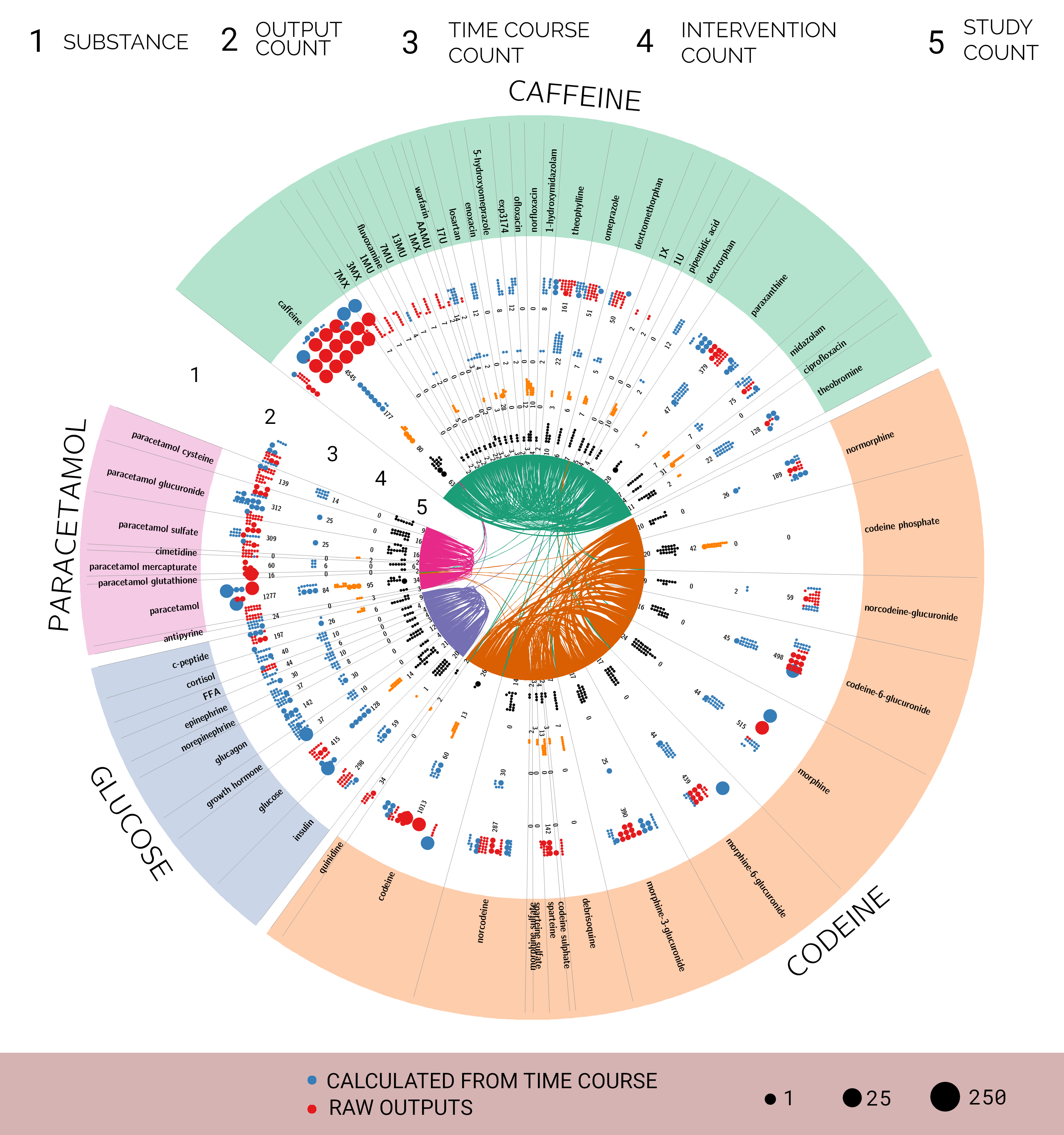
